# “Intranasal adenovirus type 5 influenza vaccination enhances milk antibody responses and passive transfer to neonatal ferrets”

**DOI:** 10.64898/2026.09.26.754702

**Authors:** Lauren Stewart Stafford, Pari Baker, Michelle Moyer, Jake Byrne, Christopher Beverly, Susanna Henry, Yixuan Bai, Yelenna Skomorovska-Prokvolit, Madelyn Cook, Ananya Goel, Francis Sun, Sean N. Tucker, Stephanie N. Langel

**Affiliations:** Department of Pathology, Center for Global Health and Diseases, Case Western Reserve University School of Medicine, Cleveland, OH, USA; Duke Human Vaccine Institute, Duke University Medical Center, Durham, NC, USA; University of Kansas School of Medicine, Kansas City, KS, USA; Office of Research Oversight, Wake Forest University School of Medicine, Winston-Salem, NC, USA; Vaxart, Inc., South San Francisco, CA, USA

## Abstract

Newborns and infants are particularly vulnerable to influenza, largely due to the immaturity of their immune systems. As such, maternal immunization is a valuable strategy to enhance influenza-neutralizing antibodies in both blood and breast milk, ensuring optimal passive immunity transfer to the infant. In our pregnant and lactating ferret maternal-neonatal model, we evaluated antibody responses to an adenovirus type 5-vectored H1N1 hemagglutinin vaccine delivered enterically, intranasally, or intramuscularly. While all three routes elicited a robust antibody response compared to saline-treated controls, intranasal vaccination significantly increased influenza-specific antibodies in dam serum, upper respiratory tract secretions, and milk, and subsequent transfer to suckling kits. Our results suggest that intranasal immunization was the most effective route for inducing functional influenza-specific antibodies in both dam serum and milk, with efficient passive transfer to suckling kits. Leveraging mucosal immune induction may enhance maternal antibody responses, transfer to offspring, and downstream neonatal protection.

## Intro

Influenza A virus is a major infectious respiratory pathogen that can lead to severe symptoms and death. The CDC estimates that in the 2024-2025 influenza season there were approximately 51 million illnesses, 710,000 hospitalizations, and 45,000 deaths in the United States^1^. Pregnant women and neonates are at increased risk of influenza infection and complications. A meta-analysis of observational influenza studies found a higher risk of influenza-associated hospitalization in pregnant women than nonpregnant women^2^. The 2024-2025 influenza season had the highest number of reported influenza-associated pediatric deaths since 2004 (excluding the 2009-2010 pandemic), with 280 reported through September 13, 2025^3,4^, 22% of which were infants less than 2 years old^5^. Mortality was highest among infants less than 6 months of age, at 11.1 deaths per million children compared with 3.8 per million overall, and these infants are ineligible for influenza vaccination on the basis of age. Of the vaccine-eligible children who died and whose vaccination status was known, 89% were not fully vaccinated^5^.

Maternal vaccination during pregnancy transfers pathogen-specific matAbs across the placenta into infant circulation and reduces risk of respiratory infection in early life. Maternal mRNA COVID-19 vaccination lowered incidence of infant hospitalizations^6–8^ and positive SARS-CoV-2 tests^9^ during the first six months of life. Maternal Tdap vaccination produces efficient transplacental transfer of anti-pertussis IgG^10^ and leads to significant infant protection through at least two months of life^10,11^. Similarly, infants born to influenza-vaccinated mothers experience fewer infections and hospitalizations during the first six months of life compared to infants of unvaccinated mothers^12–14^. Beyond transplacental transfer, maternal vaccination during pregnancy or lactation produces detectable levels of pathogen-specific IgA and IgG in breastmilk against influenza^15–17^, SARS-CoV-2^18–20^, RSV^21–23^, and pertussis^24,25^, and these breastmilk-derived antibodies have neutralizing functions *in vitro*^26–28^.

Parenteral vaccines, despite reliably inducing systemic antibodies and reducing severe disease, poorly drive local IgA responses to mucosal pathogens like influenza^29^ and SARS-CoV-2^30,31^. This is a meaningful gap, as IgA, particularly polymeric IgA, improves pathogen neutralization and has been proposed as a correlate of protection against infection^32–35^. Mucosal vaccination offers a route to this compartment, potentially inducing IgA in the maternal airway and milk while retaining systemic antibodies. A recently published clinical trial supports this, finding that an oral adenovirus type 5 (Ad5)-vectored norovirus vaccine tablet elicited significant norovirus-specific IgA in milk, as well as IgG and neutralization activity in serum^36^.

Ad5 is a non-enveloped double-stranded DNA virus that commonly infects humans and is widely used as a vaccine vector. Deletion of the E1 gene renders the virus replication-deficient and therefore unable to produce a productive infection in the host, while providing space for insertion of the transgene of interest^37^. Deletion of the E3 gene allows more space for an insertion, as well as improves immune responses to the transgene. The Ad5 vector used here expresses a TLR3 agonist as a molecular adjuvant. Ad5-vectored vaccines against several pathogens have been evaluated in pregnant and lactating humans and animals with consistent evidence of matAb transfer. All mothers vaccinated with an Ad-vectored SARS-CoV-2 vaccine (ChAdOx1-S, Ad5-nCoV, and Ad26.COV2) seroconverted, with anti-SARS-CoV-2 IgG detectable in newborn blood^38^, and maternal norovirus-Ad5 vaccination produced norovirus-specific antibodies in maternal serum and milk and in breastfeeding infants^36^. In animal models, maternal Ad5 vaccination has protected offspring against infection or death against Zika virus in mice^39^, RSV in cotton rats^40^, and porcine epidemic diarrhea virus in piglets^41^, with increased pathogen-specific IgG in neonatal serum^40^ and IgA in colostrum^41^.

Ferrets are a gold standard model for influenza studies, as they share similar sialic acid receptor expression in the upper respiratory tract (URT), symptomology, and respiratory anatomy^42–45^ to humans. Ferrets transfer little to no IgG across the placenta, and kits instead acquire maternal IgG through colostrum and milk, which crosses the permeable neonatal gut epithelium into circulation^46^. Kits therefore begin life with a pool of circulating pathogen-specific maternal IgG, as human infants do, while also receiving milk IgA at the gastrointestinal mucosa.

Using a maternal-neonatal ferret model, this study examined how the route of maternal vaccination affects influenza-specific antibody responses and their transfer to offspring.

Pregnant and lactating dams received an enteric (EN), intranasal (IN), or intramuscular (IM) Ad5-vectored hemagglutinin (HA) vaccine, and antibody responses were measured in serum, milk, and upper respiratory secretions, along with transfer to suckling kits.

## Results

### IN vaccination elicits the strongest serum antibody response

To measure influenza-specific matAb responses following vaccination, ferret dams were vaccinated with the Ad5-HA or enterically delivered saline two weeks before and two weeks after parturition. Samples were collected biweekly from Week 4 through necropsy, with a baseline sample at Week 0 (Fig. 1).

**Figure 1.**
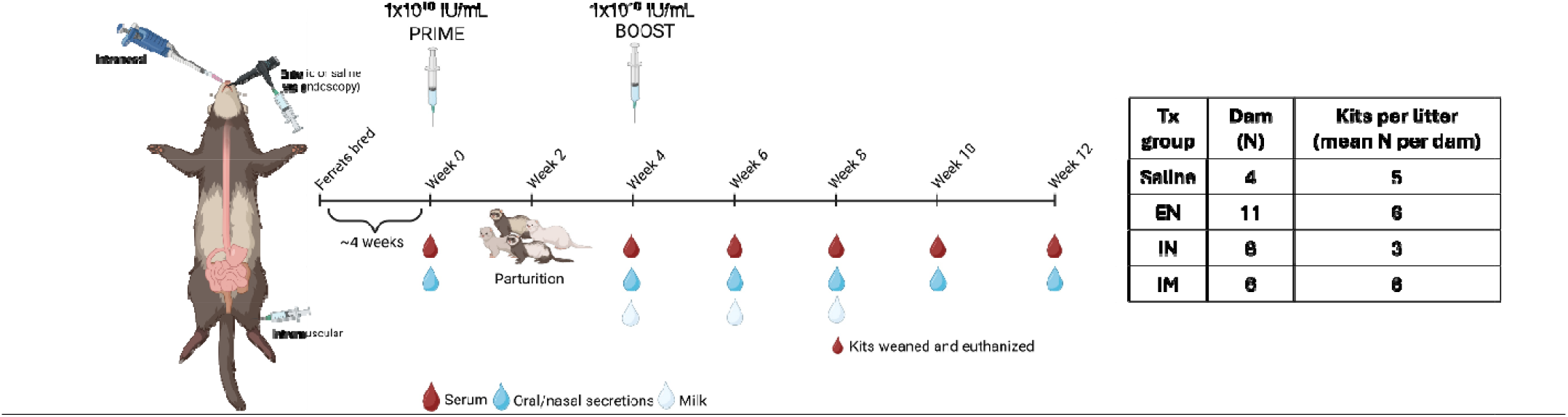
Study timeline: Pregnant ferrets were vaccinated with 1×10^10^ IU/mL Ad5-HA either enterically (via endoscopy), intranasally, or intramuscularly at 2 weeks prepartum (Week 0) and then boosted at 2 weeks postpartum, followed by sample collection every 2 weeks until necropsy (Week 12). Serum, Oral swabs, nasal washes, and milk were collected from dams at relevant timepoints, and upon weaning, kits were euthanized and serum collected. H1N1 HA-specific IgG, IgA, microneutralization, and HAI endpoint titers were measured in the collected samples to determine immunogenicity to vaccination during pregnancy and lactation by administration route. Table lists treatment group (administration route), Dam N per treatment group, and average kits per litter per treatment group.

HA-specific serum IgG titers in IN-vaccinated dams were significantly higher than those in EN- and IM-vaccinated dams at 6-, 8-, and 12-weeks post prime (Fig 2A). A significant booster effect was observed only in IN-vaccinated dams, in which titers rose from Week 4 to Week 6 (p < 0.05); titers in EN- and IM-vaccinated dams did not significantly increase over the same interval (Supplementary Fig. 1A). Ad5 vaccination increased serum IgA in all groups, but IN administration drove significantly stronger responses than EN or IM at 6-, and 12-weeks post prime (Fig. 2B). A significant boost effect was observed only in IN-vaccinated dams, in which serum IgA rose from Week 4 to Week 6 (p < 0.005), with no significant increase in EN-or IM-vaccinated dams over the same interval (Supplementary Fig 1B).

**Figure 2.**
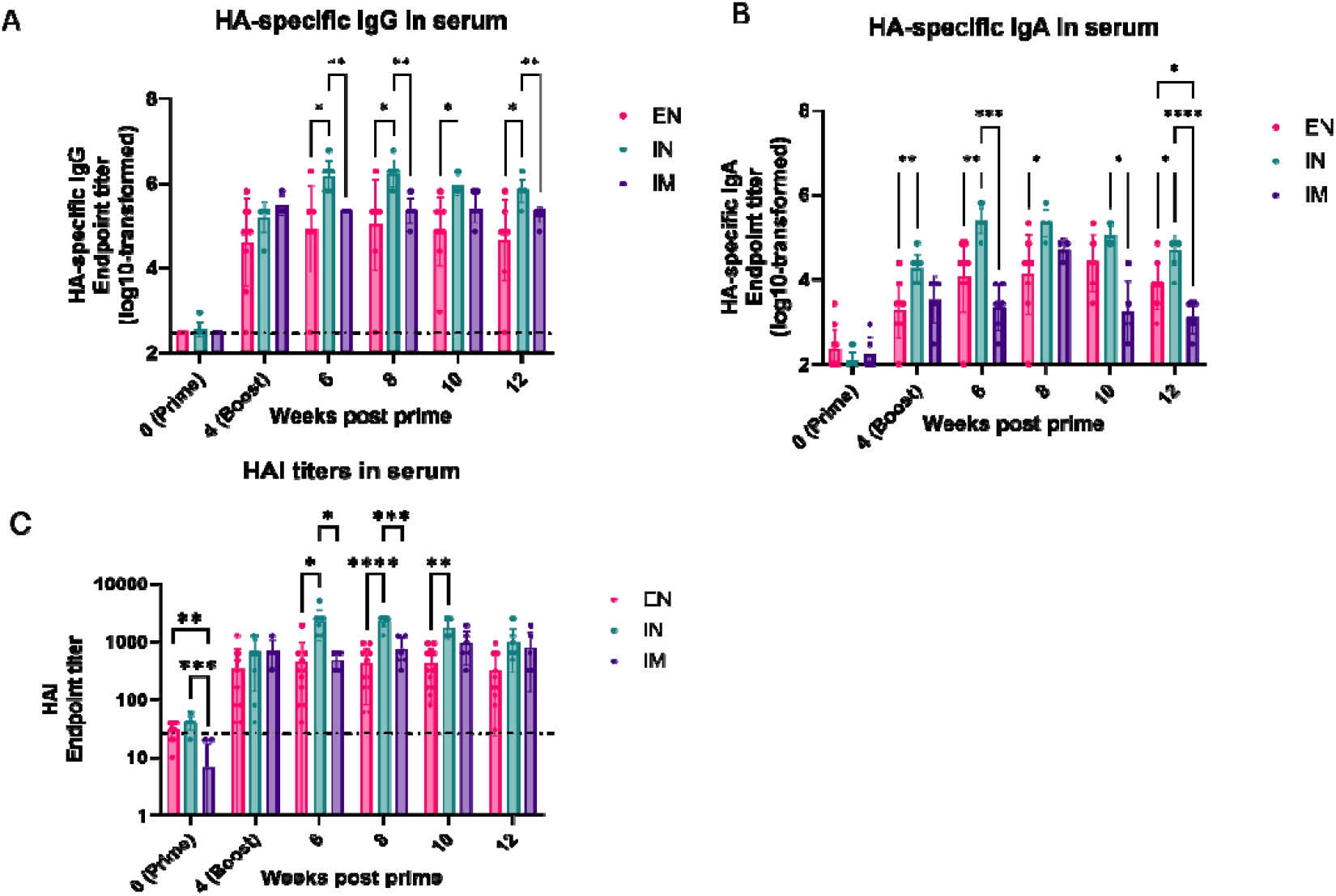
H1N1 HA-specific immunogenicity in pregnant and lactating ferret serum after Ad5-HA vaccination by enteric, intranasal, or intramuscular route. (A) HA-specific IgG endpoint titer in ferret serum up to 12 weeks after Ad5 vaccination (prime at Week 0 and boost at Week 4), (B) HA-specific IgA endpoint titer in ferret serum, and (C) Hemagglutination inhibition (HAI) endpoint titer in ferret serum. Y-axis = endpoint titers (log10-transformed for HA-specific IgG and IgA), X-axis = Weeks post prime (Week 0). Dotted line represents saline-treated endpoint titers in ferret serum. Pink = ENTERIC (EN), teal = INTRANASAL (IN), and purple = INTRAMUSCULAR (IM). Statistical analysis = Mixed-effects model with Geisser-Greenhouse correction; Tukey’s multiple comparisons test, *\*P < 0*.*05, **P < 0*.*01, ***P < 0*.*001, ****P <*□*0*.*0001*.

### IN vaccination elicits the highest serum HAI titers

To test the functional activity of HA-specific antibodies in serum, hemagglutination inhibition (HAI) and microneutralization assays were performed on receptor-destroying enzyme (RDE)-treated serum. IN vaccination led to significantly higher HAI titers than EN vaccination following the booster dose (Fig. 2C). HAI titers in serum were also significantly higher in IN-than IM-vaccinated dams at Weeks 6 (p < 0.05) and 8 post-prime (p < 0.005). A boost effect in serum HAI was observed in IN-vaccinated dams but did not reach statistical significance (p = 0.06) (Supplementary Fig. 1C). At Week 6, microneutralization titers were numerically, but not statistically, higher in IN-vaccinated dams than in EN- and IM-vaccinated dams (Supplementary Fig. 2). HAI and microneutralization titers at Week 6 were significantly positively correlated (r = 0.76, p < 0.0001).

### Route of vaccination shapes HA-specific antibody responses in milk

We next assessed antibody responses to vaccination in the milk of lactating dams. Due to the limited volume of milk obtained from some dams at individual collection time points, available post-vaccination milk samples were pooled across time points within each dam to generate a single representative sample per animal for comparison across vaccination groups. All vaccinated groups exhibited significantly higher milk HA-specific IgG responses than saline-treated animals, demonstrating a vaccine-induced antibody response in milk. Milk HA-specific IgG responses were significantly higher in IN-vaccinated dams than in EN-vaccinated dams (p = 0.03), while no significant difference was observed between IN and IM groups. Milk HA-specific IgA was significantly higher in IN-vaccinated dams than in IM-vaccinated dams (p < 0.01), whereas no significant differences were observed between the IN and EN groups (Fig. 3B).

**Figure 3.**
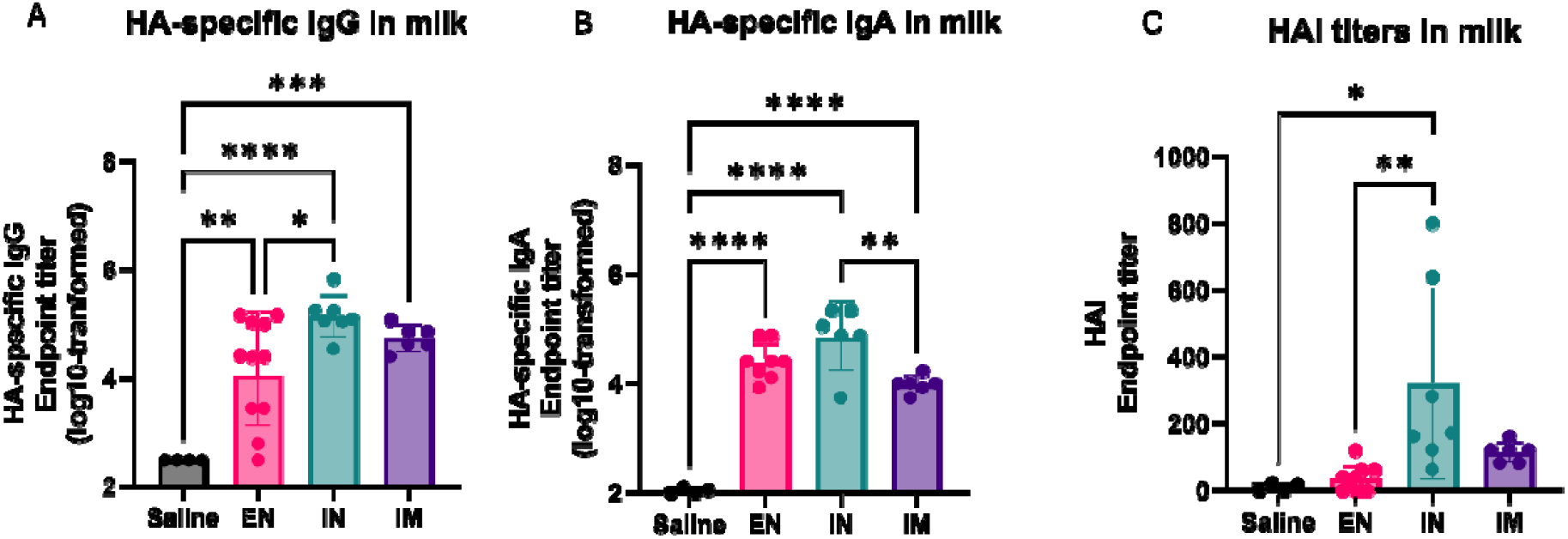
H1N1 HA-specific immunogenicity in lactating ferret milk after Ad5-HA vaccination by enteric, intranasal, or intramuscular route. (A) HA-specific IgG endpoint titer in ferret milk after Ad5 vaccination (B) HA-specific IgA endpoint titer in ferret milk, and (C) Hemagglutination inhibition (HAI) endpoint titer in ferret milk. Y-axis = endpoint titers (log10-transformed for HA-specific IgG and IgA), X-axis = Inoculation route. Gray = SALINE administration, pink = ENTERIC, teal = INTRANASAL, and purple = INTRAMUSCULAR. Statistical analysis = Ordinary one-way ANOVA with Tukey’s multiple comparison test, *\*P < 0*.*05, ** P< 0*.*01, ***P < 0*.*001, ****P <*□*0*.*0001*.

### IN vaccination elicits the highest HAI titer in milk

To assess the functional activity of vaccine-induced milk antibodies, we performed HAI assays on post-vaccination milk samples pooled from each dam. Milk HAI titers were significantly higher in IN-vaccinated dams than EN-vaccinated (p < 0.005) and saline-treated dams (p < 0.05) (Fig. 3C). HAI titers in EN- and IM-vaccinated dam milk were numerically higher compared to saline but did not reach statistical significance. Milk HA-specific IgG and HAI responses were strongly positively correlated (r = 0.8194, p < 0.0001). However, milk HA-specific IgA responses were not significantly correlated with milk HAI titers (r = 0.4, p > 0.05).

### Maternal IN vaccination drives the greatest transfer of functional antibody to suckling kits

Consistent with the maternal serum and milk responses, kits born to IN-vaccinated dams had significantly higher concentrations of HA-specific serum IgG at weaning (6–8 weeks postpartum) than kits born to EN-vaccinated dams (p < 0.005). HA-specific IgG concentrations in kits born to IN-vaccinated dams were also numerically higher than those in kits born to IM-vaccinated dams, although this difference was not statistically significant (Fig. 4A). EN-(p < 0.05) and IM-vaccination (p < 0.005) also drove IgG responses and passive transfer to kits relative to saline, though concentrations were numerically lower compared to kits from IN-vaccinated dams. Each datapoint represents the average titer for each litter. Dam milk IgG was significantly positively correlated to kit serum IgG (r = 0.73, p = 0.0001) (Fig. 4B). Kits born to IN-vaccinated dams had significantly higher HAI titers than kits born to EN-(p < 0.001) or IM-vaccinated dams (p < 0.001) (Fig. 4C). Kit serum HAI titers were also strongly positively correlated with dam milk HAI titers (r = 0.62, p < 0.005) (Fig. 4D).

**Figure 4.**
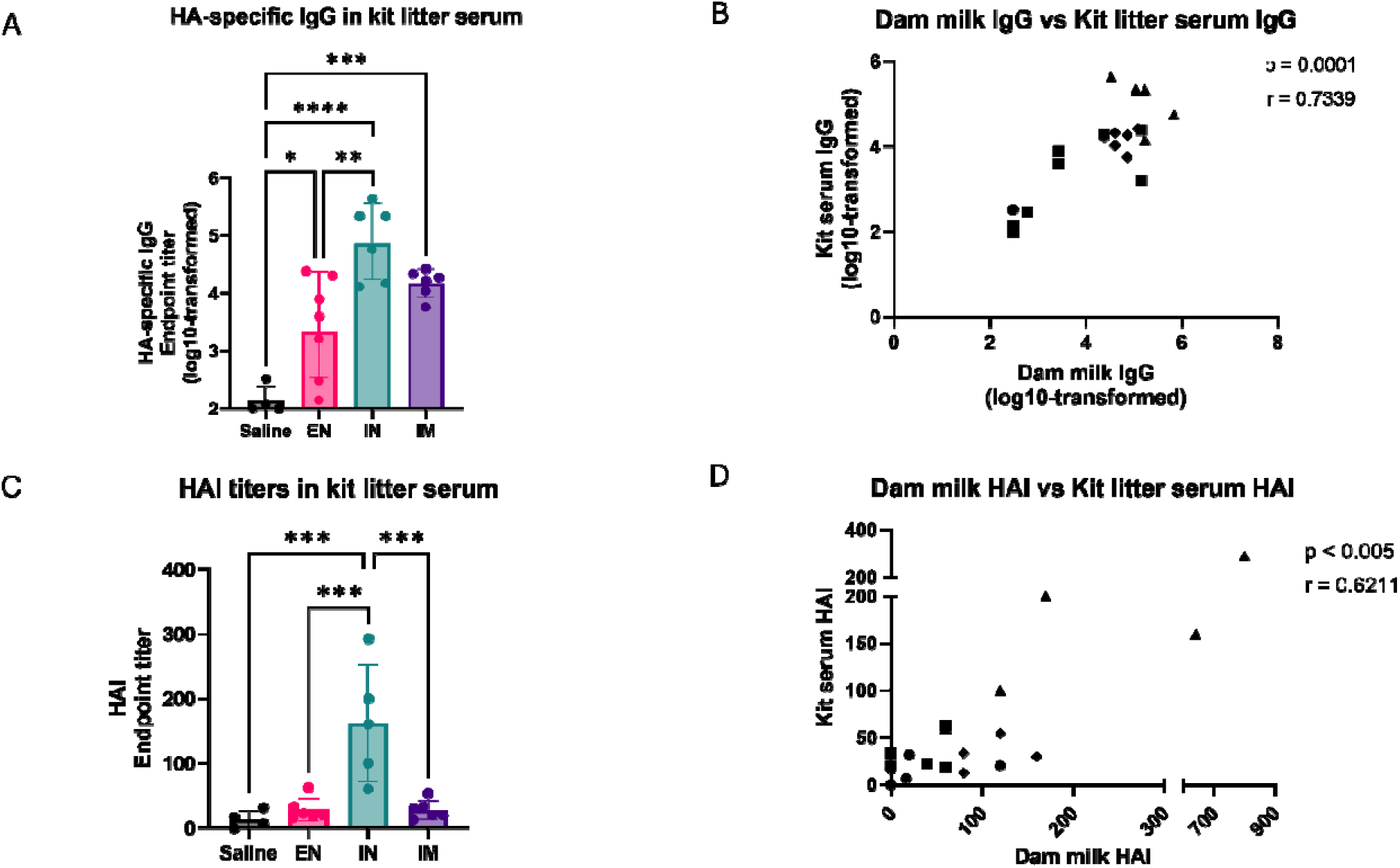
H1N1 HA-specific immunogenicity in suckling ferret kits after maternal Ad5-HA vaccination by enteric, intranasal, or intramuscular route. Each datapoint represents the average titer for each litter. (A) HA-specific IgG endpoint titer in kit litter serum after Ad5 vaccination (B) correlation of dam milk IgG endpoint titer (X-axis) to kit litter serum IgG endpoint titer (Y-axis), (C) Hemagglutination inhibition (HAI) endpoint titer in ferret kit litter serum, and (D) correlation of kit litter serum HAI titer (Y-axis) and dam milk HAI endpoint titer (X-axis). (A) and (C) Y-axis = endpoint titers (log10-transformed for HA-specific IgG and IgA), X-axis = Maternal vaccination route. Gray = SALINE (circle), pink = ENTERIC (square), teal = INTRANASAL (triangle), and purple = INTRAMUSCULAR (diamond). Statistical analysis = Ordinary one-way ANOVA with Tukey’s multiple comparisons test (A,C); Spearman correlation (B,D), *\*P < 0*.*05, **P < 0*.*01, ***P < 0*.*001, ****P <*□*0*.*0001*.

### IN leads to heightened antibody secretion in the upper respiratory tract

To measure upper respiratory tract antibody responses to vaccination, HA-specific IgG and IgA were quantified in oral swabs and nasal washes. IM vaccination elicited an early oral HA-specific IgG response peak at Week 4 post-prime that was significantly higher than responses in IN-vaccinated dams (p < 0.05) (Fig. 5A). IN-vaccinated dams exhibited statistically higher titers than EN-vaccinated dams at Week 6 only (p < 0.05) (Fig 5A). IN vaccination resulted in significantly higher HA-specific IgG in nasal secretions at Week 6 (p < 0.05) and 8 post-prime (p < 0.05) compared to EN vaccination, and at week 8 compared to IM vaccination (p < 0.05) (Fig. 5B). Only IN vaccination elicited a boost effect in IgG levels in the URT (p < 0.01 in oral; p < 0.05 in nasal), not observed after in EN or IM vaccination (Supplementary Fig. 3A,B). Compared to EN- and IM-vaccinated dams, IN-vaccinated dams had higher IgA in oral swabs at Week 6 (p < 0.05) (Fig. 5C). HA-specific IgA in nasal secretions were numerically higher in IN-vaccinated dams but did not differ significantly from EN- and IM-vaccinated dams until Week 12 (IM only) (Fig. 5D). No boost effect was observed in URT HA-specific IgA.

**Figure 5.**
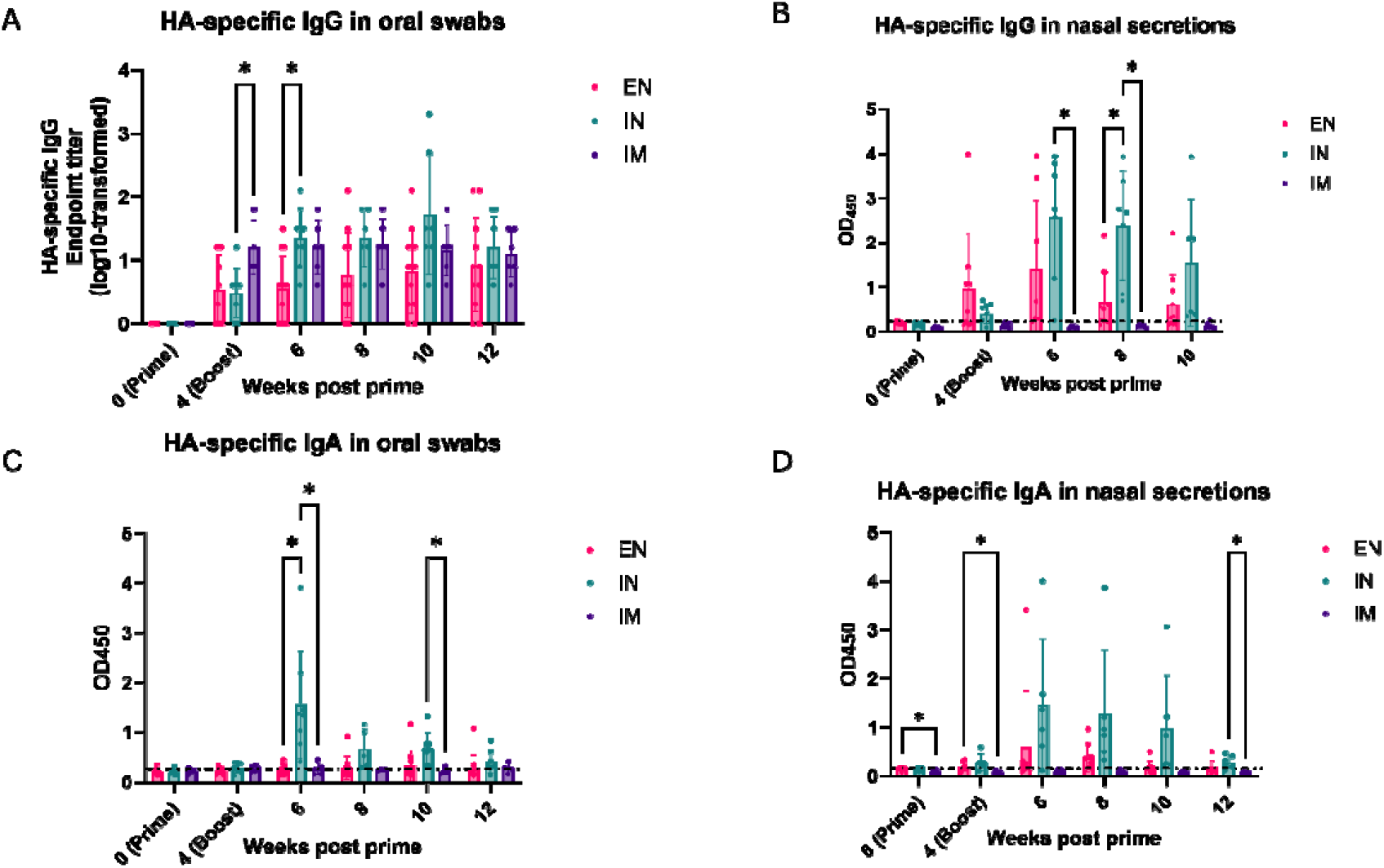
H1N1 HA-specific immunogenicity in pregnant and lactating ferret upper respiratory tract after Ad5-HA vaccination by enteric, intranasal, or intramuscular route. (A) HA-specific IgG endpoint titer in ferret oral swabs up to 12 weeks after Ad5 vaccination (prime at Week 0 and boost at Week 4), (B) HA-specific IgG OD450 in ferret nasal secretions, (C) HA-specific IgA OD450 in oral swabs, and (D) HA-specific IgA OD450 in nasal secretions. (A) Y-axis = endpoint titers (log10-transformed for HA-specific IgG), (B) Y-axis = OD450, X-axis = Weeks post prime (Week 0). Dotted line represents saline-treated endpoint titers in ferret URT samples. Pink = ENTERIC (EN), teal = INTRANASAL (IN), and purple = INTRAMUSCULAR (IM). Statistical analysis = Mixed-effects model with Geisser-Gree house correction; Tukey’s multiple comparisons test, *\*P < 0*.*05, **P < 0*.*01, ***P < 0*.*001, ****P <*□*0*.*0001*.

### Pregnancy and lactation do not alter the antibody response to EN vaccination

Given the dynamic physiological and immunological changes that occur in the gastrointestinal tract during pregnancy^47,48^, we sought to determine whether pregnancy altered antibody responses to enteric vaccination. Aged-matched, non-pregnant or lactating female ferrets were vaccinated EN with the same dose of Ad5-HA and sampled over the same schedule. No significant differences in serum HA-specific IgG or IgA responses were observed between pregnant and lactating dams and non-pregnant, non-lactating animals (Supplementary Fig. 4), oral swabs, or nasal secretions (Supplementary Fig. 5) suggesting that pregnancy did not substantially alter the antibody response to EN vaccination.

### Route-dependent anti-vector antibody responses following Ad5-HA vaccination

One potential limitation to adenoviral vector vaccines is the induction of anti-vector antibodies that may blunt immune responses to the encoded antigen following subsequent doses. We therefore measured anti-Ad5 IgG responses in a subset of vaccinated dams. Anti-Ad5 IgG was undetectable in animals prior to vaccination (Week 0). At two weeks post-boost (Week 6), little to no anti-Ad5 reactivity was detected in serum from EN-vaccinated dams, whereas anti-Ad5 reactivity significantly increased in IN- and IM-vaccinated dams compared to saline-treated controls and baseline titers (Week 0) (Supplementary Fig. 6). Despite this, IN-vaccinated dams mounted significant HA-specific IgG responses post-boost (Fig. 2A), indicating that anti-Ad5 immunity did not preclude antibody responses to the vaccine-encoded HA antigen.

## Discussion

Maternal immunization protects both the pregnant individual and, through passively transferred antibodies, the infant^49,50^. How that protection is distributed depends on route: parenteral vaccination can increase systemic IgG but poorly induces mucosal IgA^29–31^, leaving the respiratory and mammary compartments comparatively underserved. In this study, we examined how the route of administration of an Ad5-HA influenza vaccine affects systemic and mucosal antibody responses in a maternal-neonatal ferret model. IN vaccination was the only route to drive a robust HA-specific IgA response, significantly exceeding IM and EN in serum and in oral secretions, with the same pattern in nasal secretions and milk. While IN-vaccinated dams had significantly higher serum and milk HA-specific IgG than EN-vaccinated dams, they did not differ significantly from IM-vaccinated dams in either compartment. Functionally, however, the IN advantage in ferrets was consistent. IN-vaccinated dams had the highest serum HAI and microneutralization titers at six-weeks post-boost, the highest milk HAI titers, and their kits reached weaning with significantly higher serum HAI than kits from either of the other administration routes. The route of maternal vaccination in ferrets therefore shaped not only the magnitude of the maternal response but the functional quality of antibody the neonate received.

The route-dependent IgA responses we observed are consistent with the common mucosal immune system, a framework established in rodent and ruminant models in which antigen encounter at one mucosal inductive site generates antigen-specific IgA at distant effector sites, including the lactating mammary gland^51,52^. In these systems, antibody-secreting cells mostly from the intestine are recruited to the mammary gland through CCL28 produced by mammary epithelium, which binds CCR10 on IgA plasmablasts^53–55^. The relevance of this pathway extends to humans, where oral tablet immunization of lactating participants with an Ad5-vectored, TLR3-adjuvanted norovirus vaccine, the same platform used in the present study, produced norovirus-specific IgA in both serum and milk^36^.

Given this, we anticipated that EN vaccination would be the most effective route for driving HA-specific IgA into ferret milk. In our study, EN vaccination elicited HA-specific IgA in milk relative to saline-treated animals. IN delivery, by contrast, elicited the highest HA-specific IgA across serum, upper respiratory secretions, and milk and was also associated with the greatest transfer of functional antibody to kits. The presence of HA-specific IgA in milk following respiratory antigen delivery is consistent with mucosal exposure at one site generating antibody in a distant effector compartment. Because our data are limited to antibody endpoints, we cannot determine whether this reflects trafficking of HA-specific plasmablasts or plasma cells to the mammary gland, or transfer of systemic antibody into milk. It is also possible that enteric immunization of ferrets is inefficient compared to human tablet delivery.

Live attenuated influenza vaccines (LAIV) delivered intranasally induce local IgA secretion^56^, diverse T cell responses^57^, and early activation of anti-influenza immune pathways in milk^58^.

Inactivated influenza vaccine (IIV), given intramuscularly, reliably raises systemic IgG but induces mucosal IgA poorly, consistent with the weak IgA responses we observed across serum, milk, and the upper respiratory tract following IM delivery^29–31^.

Ferret kits, unlike human infants, acquire circulating maternal IgG almost exclusively through colostrum and milk rather than transplacentally^59^. Consistent with the maternal serum and milk data, kits born to IN-vaccinated dams had the highest concentrations of HA-specific IgG in serum at weaning, and significantly higher HAI titers than kits born to either EN-or IM-vaccinated dams. Dam milk IgG was strongly correlated with kit serum IgG, and dam milk HAI with kit serum HAI, indicating that both the magnitude and functional quality of the maternal milk response shape what the neonate receives. Applying the 1:40 HAI titer conventionally associated with 50% protection in adult humans as a benchmark^60,61^, all kits from IN-vaccinated dams reached or exceeded this threshold at weaning compared with 41% and 37% of kits from EN- and IM-vaccinated dams, respectively.

Milk IgG rather than milk IgA tracked with functional activity in both compartments, which is expected given the IgG-dominant immunoglobulin profile of ferret^46,62,63^ and closely related carnivore milk^64,65^. The route-dependent milk IgA responses described above likely reflect mucosal induction in the dam rather than a contribution to systemic transfer to kits. In humans, where milk is approximately 90% IgA^49,66^, the route-dependent induction of milk IgA we observed would be expected to contribute more to infant protection than in the ferret model.

Mucosal immunization with this vector platform has already been shown to elicit pathogen-specific IgA in the milk of lactating participants^36^.

The route of administration also shaped the response to the vector itself. Anti-Ad5 IgG was undetectable in tested animals prior to vaccination and remained near-undetectable in EN-vaccinated dams, whereas IN and IM vaccination elicited readily measurable responses. This may reflect less vector reaching or transducing cells at the enteric site, or a lower antibody response to comparable exposure, as the gut mucosa is a tolerogenic environment in which responses to orally encountered antigen are often attenuated^67^. Formulation and delivery may also contribute, as the human oral product is administered as an enterically coated tablet that releases in the small intestine^68,69^, whereas in our study the vector was delivered endoscopically as a liquid, which bypasses gastric transit but may differ in release site, volume, and residence time. Quantifying vector genome copies or transgene expression in intestinal tissue would distinguish reduced delivery from an attenuated response.

Both IN- and IM-vaccinated dams developed anti-Ad5 IgG, yet only IN-vaccinated dams mounted a significant HA-specific boost response. Whether this reflects anti-vector interference or route-intrinsic differences in immunogenicity cannot be answered from these data. This anti-vector and anti-HA boost is relevant for translation, as anti-Ad5 seroprevalence ranges from approximately 40% in the United States and Europe to as high as 75% in parts of Asia^70,71^. Pre-existing anti-Ad5 immunity attenuates but does not abolish responses to Ad5-vectored vaccines which retained 48% effectiveness against COVID-19 infection in populations with substantial seroprevalence^72–74^, particularly when delivered intranasally^75,76^. Our animals were Ad5-naive at prime, however, and responses in a seroprevalent population may be attenuated relative to those reported here.

Kits were not challenged with influenza, so kit serum HAI titers represent a correlate of immunity rather than demonstrated protection. The 1:40 threshold applied here derives from adult human data and has not been validated in ferrets or neonates, who may require higher titers for protection. The scarcity of validated ferret reagents constrained our analysis to antibody endpoints, precluding direct assessment of the plasmablast trafficking that would establish the mucosal pathway we propose. Limited milk volume at individual timepoints also required pooling within each dam, preventing longitudinal analysis of the milk antibody response.

This preclinical study shows that maternal IN Ad5-HA vaccination leads to significant influenza-specific IgA, IgG, and inhibitory effects in both serum and milk. This maternal immune response is then passively transferred to suckling kits. Due to the efficient transfer of pathogen-specific matAbs during gestation and/or lactation, maternal vaccination is an attractive public health measure. Our results suggest that local, IN immune activation may be an effective strategy for generating anti-influenza responses that protect mothers and passively, their infants.

## Materials and Methods Study design

This preclinical study used a maternal-neonatal ferret model to evaluate the magnitude and functional activity of antibody responses following maternal vaccination with an Ad5-vectored influenza HA vaccine delivered by enteric, intranasal, or intramuscular route.

### Biosafety

The recombinant adenoviral vector vaccine is derived from adenovirus type 5 (Ad5) and has deletions in the E1 and E3 regions, rendering the virus replication-deficient. Adenoviral vectors are nonetheless handled as a potential hazard, and extra precautions were taken during immunization. Personnel wore a head cover, N95 respirator and face shield, disposable lab gown, shoe covers, and two pairs of gloves. Immunizations were carried out within a class II biological safety cabinet, and syringes and tubes containing the vaccine were decontaminated before disposed into biohazardous waste. The animal housing room was considered potentially infectious for 72 hours following vaccination, and the same personal protective equipment was required during that period.

### Animal care and welfare

Upon arrival, pregnant ferrets were individually housed in stainless-steel cages (Allentown Inc.). Ferrets were provided with food and freshwater *ad libitum*. At parturition, ferret dams and kits remained unperturbed and remained housed together until weaning at approximately 6-8 weeks postpartum. Animals received daily veterinary monitoring and care, and examinations were performed by trained veterinary staff prior to all procedures.

### Cells and viruses

MDCK cells (propagation) and MDCK-SIAT1 cells (microneutralization), provided by Dr. Georgia Tomaras (Duke University, Durham, NC), were maintained in Dulbecco’s Modified Eagle Medium (DMEM) with GlutaMAX (ThermoFisher) supplemented with 10% fetal bovine serum (FBS) and 1X penicillin-streptomycin and incubated with 5% CO_2_ at 37°C. Viral assays were performed using A/California/07/2009 H1N1 obtained from the Duke Human Vaccine Institute and propagated in MDCK cells in Minimum Essential Medium (MEM) with Earle’s Salts and L-glutamine, supplemented with 1X penicillin-streptomycin, 0.13% bovine serum albumin (BSA), 1mM sodium pyruvate, 1X non-essential amino acids (NEAA), 10mM HEPES, and 3mg/mL TPCK-trypsin. Turkey red blood cells (RBCs, Innovative Research) were supplied as a 5% suspension, then washed and diluted to 0.5% in PBS for hemagglutination inhibition assays.

### Vaccine preparation

VXA-A1.1 is an E1/E3-deleted, replication-incompetent, adenovirus type 5 vaccine designed as a vaccine for prevention of H1N1 influenza, was provided by Vaxart, Inc. (South San Francisco, CA). The vaccine vector encodes a 1.7 kb hemagglutinin (HA) gene from the A/California/04/2009 H1N1 influenza strain. The HA gene is codon-optimized for expression in mammalian cells and is expressed using a human cytomegalovirus (CMV) intermediate early region (hCMVIE) enhancer/promoter with a bovine growth hormone (BGH) polyadenylation (pA) signal. This expression cassette also includes the first intron of human ß-globin to enhance transgene expression. In addition to the transgene cassette, a second hCMVie promoter is used to express a double-stranded RNA sequence that acts as a TLR3 agonist adjuvant. VXA-A1.1 was delivered at a total dose of 1 × 10^10^ IU by each route. Enteric administration was by endoscopy in a 10mL total volume, IN administration as 0.5 mL per nare, and intramuscular administration as 0.5 mL per thigh.

### Ferret vaccination

Pregnant female ferrets were sourced from Marshall BioResources at approximately 3 weeks prepartum. Animals were vaccinated at 2 weeks prepartum and a booster dose of the same vaccine at 2 weeks postpartum, with dose and route as described above. Control animals received sterile saline by the enteric route on the same schedule. All ferret experiments were approved by the Institutional Animal Care and Use Committees (IACUC) at Duke University (A225-20-11) and Case Western Reserve University (2022-0090).

### Sample collection

Sample collections were performed at weeks 0 (2 weeks prepartum), 4 (2 weeks postpartum), 6, 8, 10, and 12 (necropsy). Animals were not handled at week 2 to avoid disturbance around parturition. Sample collections were performed under sedation with Midazolam (0.5 mg/kg) and Dexmedotomidine (0.025mg/0.05mL) delivered intramuscularly, with anesthesia maintained using 2% isoflurane and an oxygen flow rate of 2L per minute via nose cone. Blood was collected via venipuncture from the cranial vena cava into additive-free serum collection tubes and left undisturbed for 30 minutes at room temperature to allow blood clot formation. Oral swabs were placed into 1 mL of PBS. Nasal washes were performed by instilling 0.5mL of PBS into each nare and allowing the liquid to drip back into the collection tube. Milk was expressed from all available active teats (∼0.2-1mL total volume) into a microcentrifuge tube. After weaning, blood was collected from kits as described above, after which kits were euthanized under anesthesia with pentobarbital sodium (195mg/0.5mL) administered via the cranial vena cava. Dams were euthanized by the same method at necropsy (390mg/mL).

### Sample processing

The blood clot was removed by centrifuging at 900 × g for 10 minutes at 4°C, and the supernatant transferred and centrifuged again at 2000 × g for 15 minutes at 4°C. Serum was stored at -80°C for further assays. Oral and nasal swab eluate samples were centrifuged at 4000 × g for 10 minutes at 4°C, and supernatants stored at -80°C. Milk was diluted 1:1 in PBS and centrifuged at 1200 × g for 15 minutes. The aqueous layer between the cell pellet and the fat layer was collected by pipette, transferred to a SpinX centrifuge tube filter (0.22 µm cellulose acetate, Corning, Corning, NY), and centrifuged at 15000 × g for 15 minutes to remove residual lipid. The final filtrate was stored at -80°C.

### HA-specific ELISA

384-well plates were coated overnight at 4°C with 2 μg/mL A/California/07/2009 HA antigen (eEnzyme) in coating buffer (SeraCare). Plates were washed once and blocked with SuperBlock (4% whey, 15% goat serum, and 0.5% Tween-20 in PBS) for 1-2 hours at room temperature.

Samples were serially diluted according to sample type, with serum and milk diluted 3-fold from a starting dilution of 1:300, oral swab eluates diluted 2-fold from 1:1, and nasal washes assayed undiluted. Plates were washed once, samples added and incubated at room temperature for 1-2 hours. Plates were washed four times, and secondary antibody added: goat anti-ferret IgG-HRP (Bethyl Cat #A140-108P) at 1:10,000, or goat anti-dog IgA-HRP (Bethyl Cat #A40-104P) at 1:1000. Plates were incubated at room temperature for 1 hour, washed four times, and developed with TMB substrate (SeraCare) for 2-3 minutes before addition of an equal volume of stop solution. Absorbance was read at 450 nm on a Cytation 1 Cell Imaging Multi-Mode Reader (Agilent BioTek). HA-specific IgG endpoint titers were determined as the reciprocal of the highest sample dilution with an absorbance exceeding the mean plus three standard deviations of the saline-treated control samples at all timepoints. Nasal wash and oral swab HA-specific IgA and nasal wash IgG responses did not reach titratable levels and are reported as absorbance at 450 nm.

### Anti-Ad5 ELISA

Ad5 vector vaccine was heat-inactivated for 20 minutes at 60°C and used to coat a 384-well plate overnight at 2 μg/mL inactivated stock in coating buffer (SeraCare). Serum samples were diluted 1:10 and ran in 3-fold serial dilutions. The assay was otherwise performed as described above.

### Meso Scale Discovery (MSD)

96-well One-Spot Sector plates (Meso Scale Discovery) were coated overnight at 4°C with 6.6 ng/μL A/California/07/2009 HA antigen (eEnzyme) in PBS. Plates were washed three times and blocked for 1 hour in 3% whey in PBS with 0.1% Tween-20. After three washes, serum and milk samples were added at a starting dilution 1:300 with 3-fold serial dilutions and incubated with shaking at room temperature for 2 hours. Plates were washed three times, and sulfotagged secondary antibody was added: anti-dog IgA (BioRad) at 1:1,000 for serum and anti-ferret IgA (Bethyl Cat# A140-113A, discontinued) at 1:2,000 for milk. Plates were incubated shaking at room temperature for 1 hour, washed three times, and Read Buffer T 4X (Meso Scale Discovery) was added at a 1:2 dilution. Plates were immediately read on an MSD (1300 MESO QuikPlex SQ 120MM) plate reader. HA-specific IgA endpoint titers were defined as the reciprocal of the highest sample dilution exceeding the mean plus three standard deviations of saline-treated control samples across all timepoints.

### Hemagglutination inhibition assay (HAI)

Serum samples treated with receptor-destroying enzyme (Hardy Diagnostics) were diluted 1:10 and serially diluted two-fold in PBS in 96-well V-bottom plates to 25uL total volume/well. 25uL of undiluted RDE-treated serum was placed in column 1 with 25uL PBS to act as a serum-only control to account for serum-RBC reaction. 25uL of A/California/07/2009 H1N1 virus, diluted in PBS to 8 HAU/50uL, was added to columns 2-11 and incubated at room temperature for 30 minutes. Turkey RBCs (0.5%, 50 µL) were then added to all wells and incubated at room temperature for 1 hour to allow RBC precipitation. Titers were scored manually by the presence of an RBC button, indicating inhibition of hemagglutination, and defined as the reciprocal of the highest dilution showing no agglutination.

### Microneutralization

RDE-treated serum samples were diluted 1:10 in virus diluent (DMEM with 1X penicillin-streptomycin, 1% BSA, 10nM HEPES, and 1 μg/mL TPCK-trypsin), followed by a 2-fold serial dilution in flat-bottom 96-well cell culture-treated plates. A/California/07/2009 H1N1 was diluted in virus diluent to 100 TCID_50_ per 50 μL, added to the plate, and incubated for 1 hour at 37°C. MDCK-SIAT1 cells were then added at 20,000 cells per 100 μl and incubated for 18-22 hours at 37°C. Media was aspirated into 10% bleach solution, wells were washed with PBS, and cells were fixed with cold 80% acetone in PBS for 10 minutes. Plates were washed three times in wash buffer (PBS with 0.1% Tween-20), and primary antibody (anti-Influenza A NP monoclonal antibodies, clones A1 (Millipore Sigma Cat# MAB8257) and A3 (Millipore Sigma Cat# MAB8258) added at 1:4,000 for 1 hour at room temperature. Plates were washed three times, and goat anti-mouse IgG-HRP (ThermoFisher Cat# 31430) added at 1:4,000 for 1 hour at room temperature. Plates were washed four times with wash buffer and once with water, developed with TMB substrate (SeraCare) for 2-5 minutes, and the enzymatic reaction was stopped by adding an equal volume of stop solution. Absorbance was immediately read on a Cytation 1 Cell Imaging Multi-Mode Reader (Agilent Biotek). Neutralization titers were defined as the reciprocal of the highest dilution at which absorbance fell below the 50% infectivity cutoff, calculated as the mean absorbance of virus-only wells plus the mean of cells-only wells, divided by two.

### Statistical analysis

Unless otherwise noted, HA-specific antibody titers were log_10_-transformed prior to analysis and figure presentation. Ordinary one-way ANOVA with Tukey’s correction was used to compare the effects of administration route (EN, IN, or IM) in kit serum and milk. Mixed effects two-way ANOVA with the Geisser-Greenhouse correction was used to assess differences across route and timepoint in serum, oral swab, and nasal wash samples, with animals treated as matched across timepoints. Correlations between dam milk and kit serum measures were assessed by Spearman correlation,. All analyses were performed using GraphPad Prism version 10.3, and p values ≤ 0.05 were considered statistically significant.

## Acknowledgments

This work was funded by the Bill & Melinda Gates Foundation (INV-22595D). We thank the staff of the Animal Resource Center at Case Western Reserve University and the Division of Laboratory Animal Resources at Duke University for veterinary care and animal husbandry support.

## Contributions

Conceptualization: S.N.L. and S.T. Investigation: L.S.S., P.B., M.M., J.B., C.B., S.H., Y.B., Y.S-P., M.C., A.G., F.S., S.N.L. Data Curation: L.S.S., P.B., J.B., C.B., S.H., M.C., A.G. Formal Analysis: L.S.S., P.B., J.B., C.B., S.H. Visualization: L.X.S. and S.N.L. Resources: S.N.L. and S.T. Writing - Original Draft: L.S.S. and S.N.L. Writing - Review and Editing: L.S.S., P.B., M.M., J.B., C.B., S.H., Y.B., Y.S-P., M.C., A.G., F.S., S.T., S.N.L.

Corresponding authors: Lauren Stewart Stafford and Stephanie N. Langel

## Ethics declaration

S.T. is an employee and shareholder of Vaxart, Inc. All other authors declare no competing interests.

