## Supplementary Figures for "“Intranasal adenovirus type 5 influenza vaccination enhances milk antibody responses and passive transfer to neonatal ferrets”": Stafford_2026_figures.pptx

### Slide 1
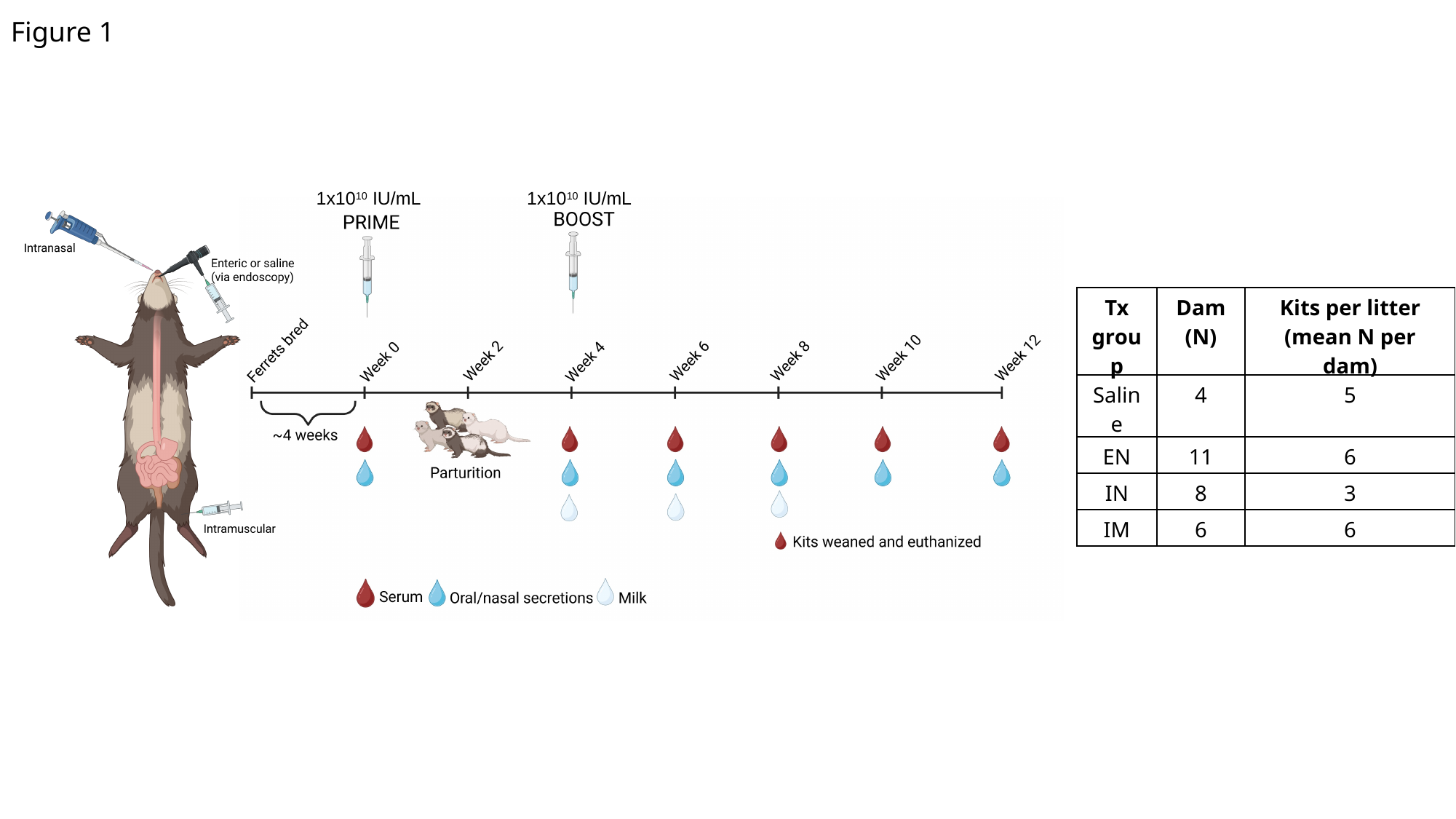

Figure 1
1x10^10^ IU/mL
1x10^10^ IU/mL
| Tx group | Dam (N) | Kits per litter (mean N per dam) |
| --- | --- | --- |
| Saline | 4 | 5 |
| EN | 11 | 6 |
| IN | 8 | 3 |
| IM | 6 | 6 |

### Slide 2
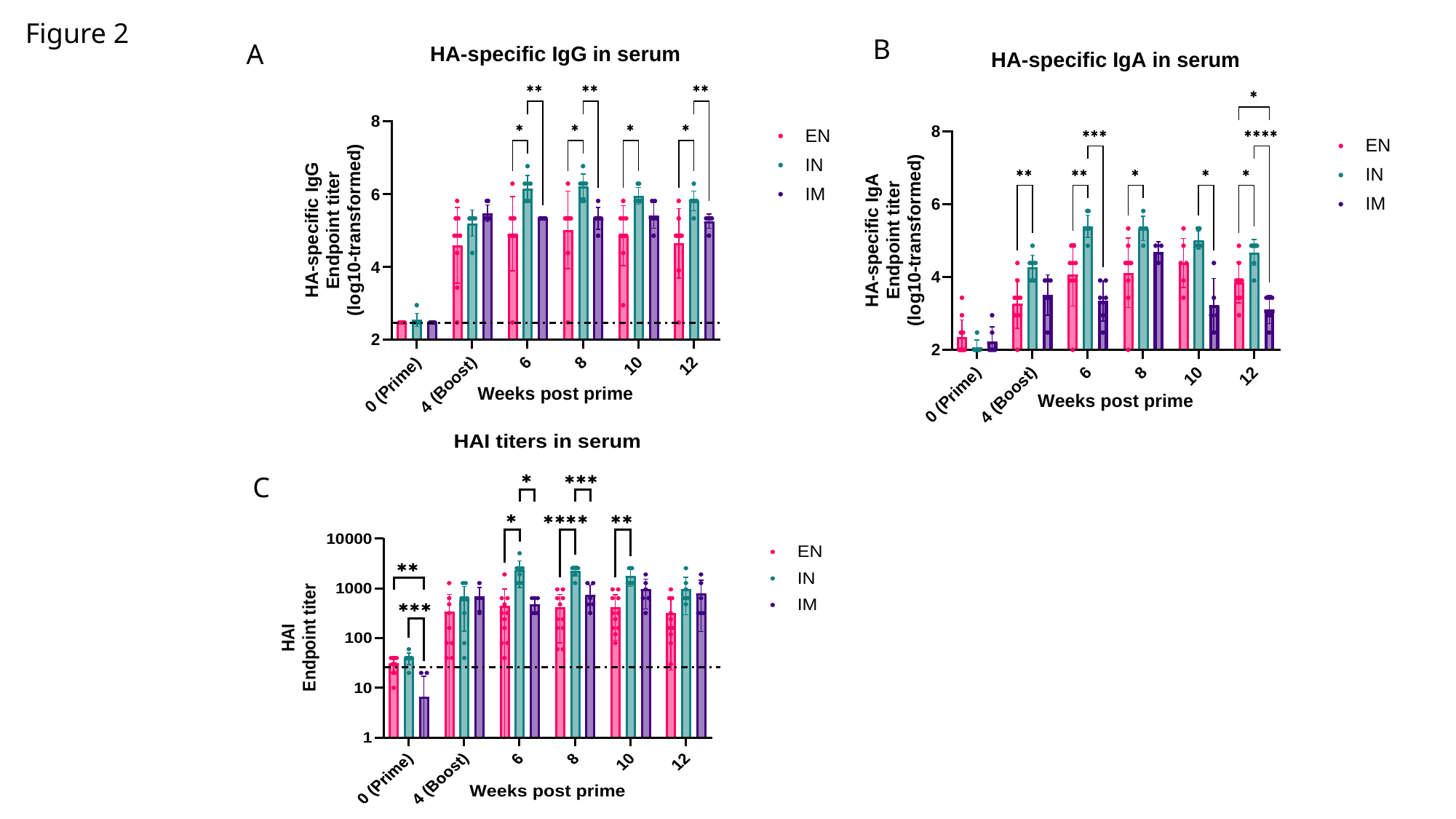

Figure 2
B
A
C

### Slide 3
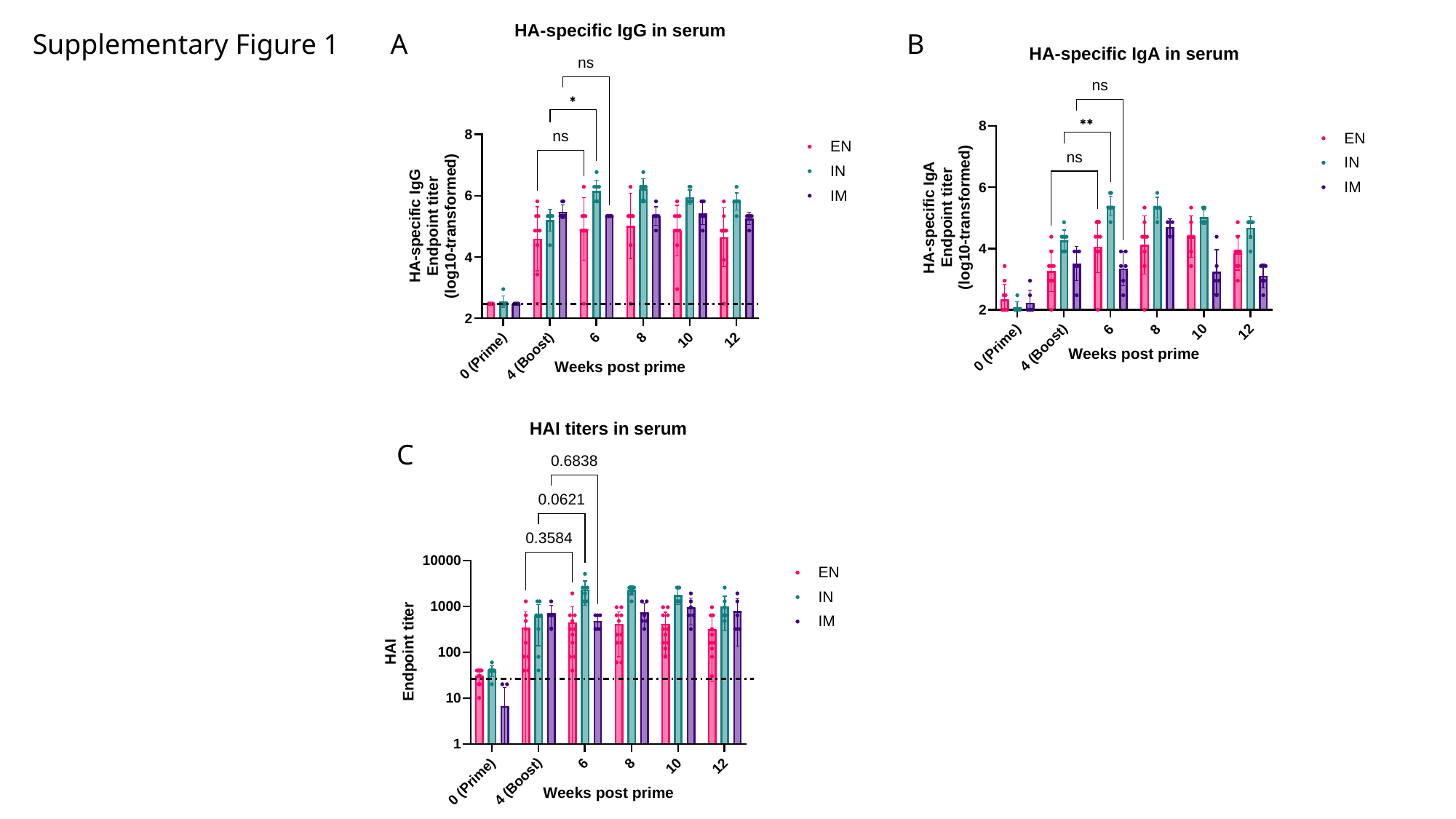

Supplementary Figure 1
A
B
C

### Slide 4
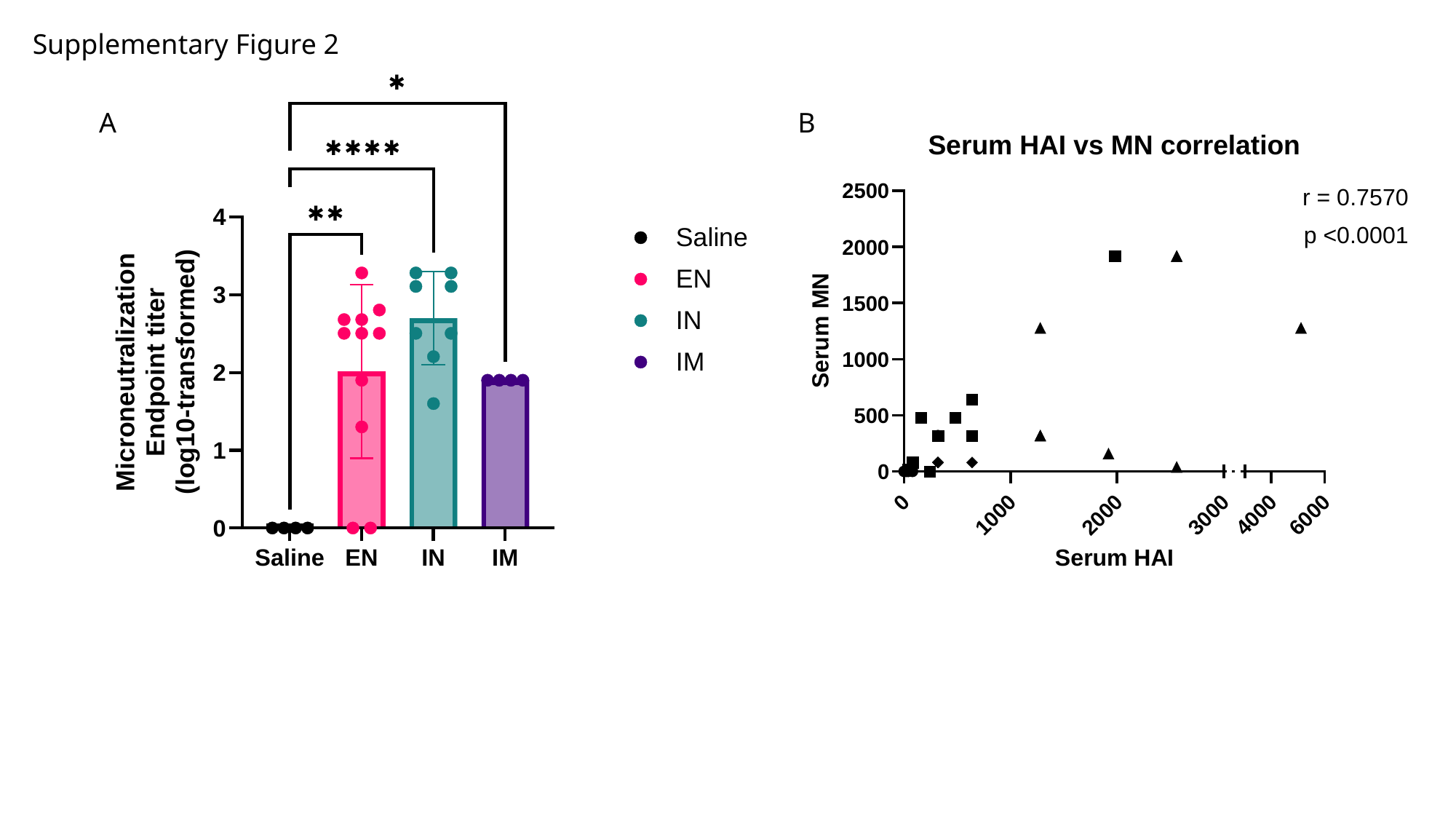

Supplementary Figure 2
A
B

### Slide 5
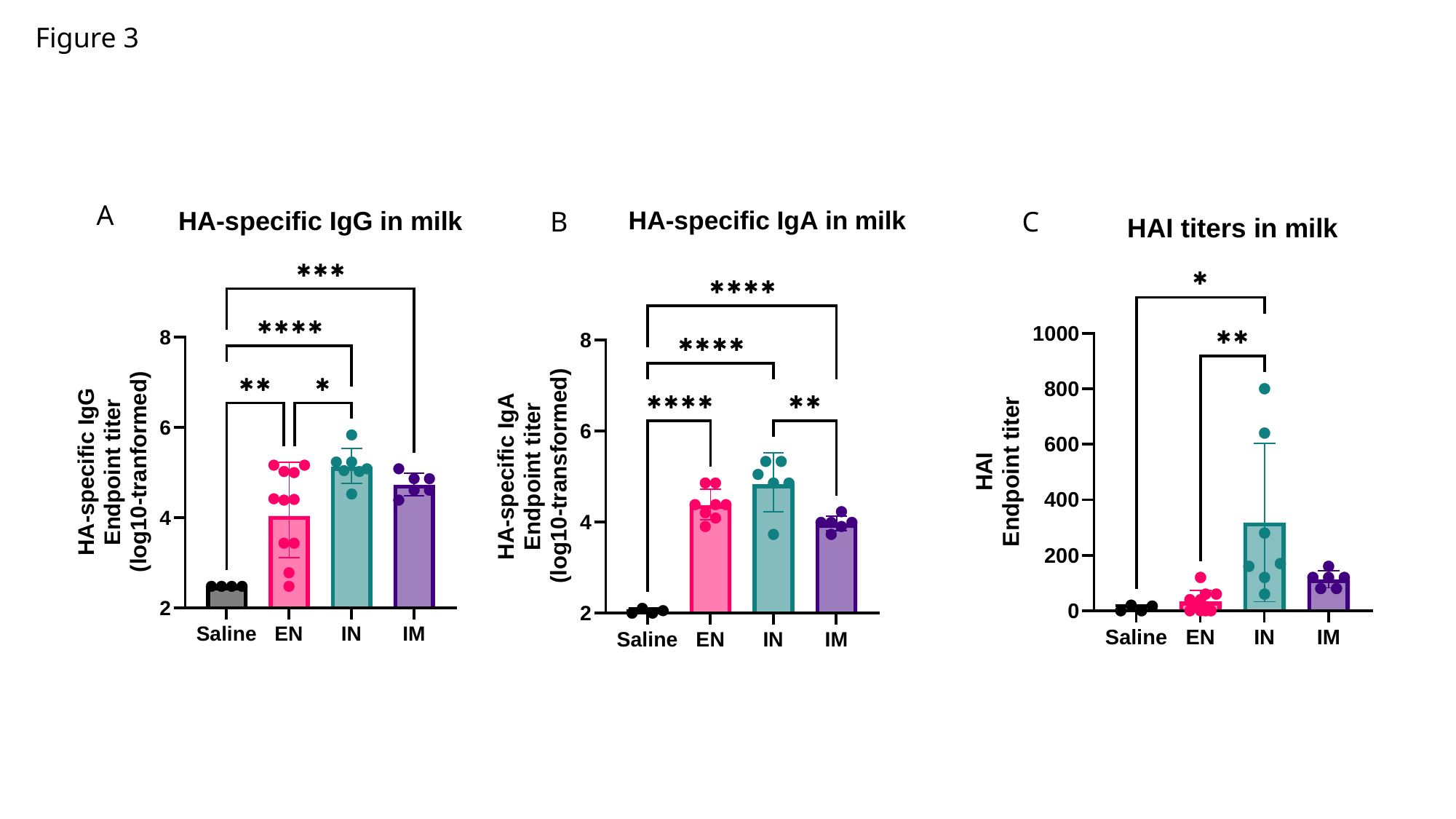

Figure 3
A
B
C

### Slide 6
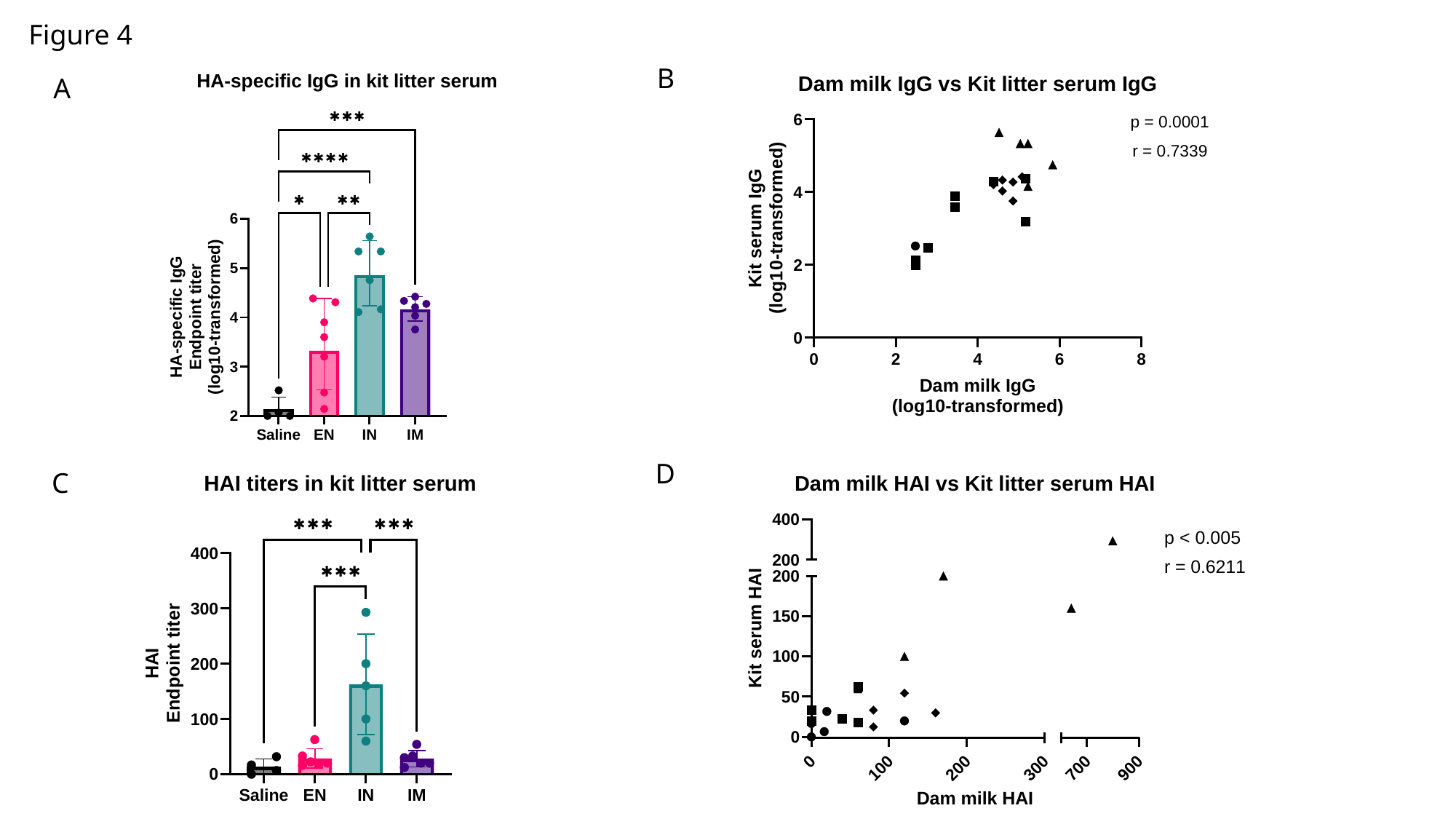

Figure 4
B
A
D
C

### Slide 7
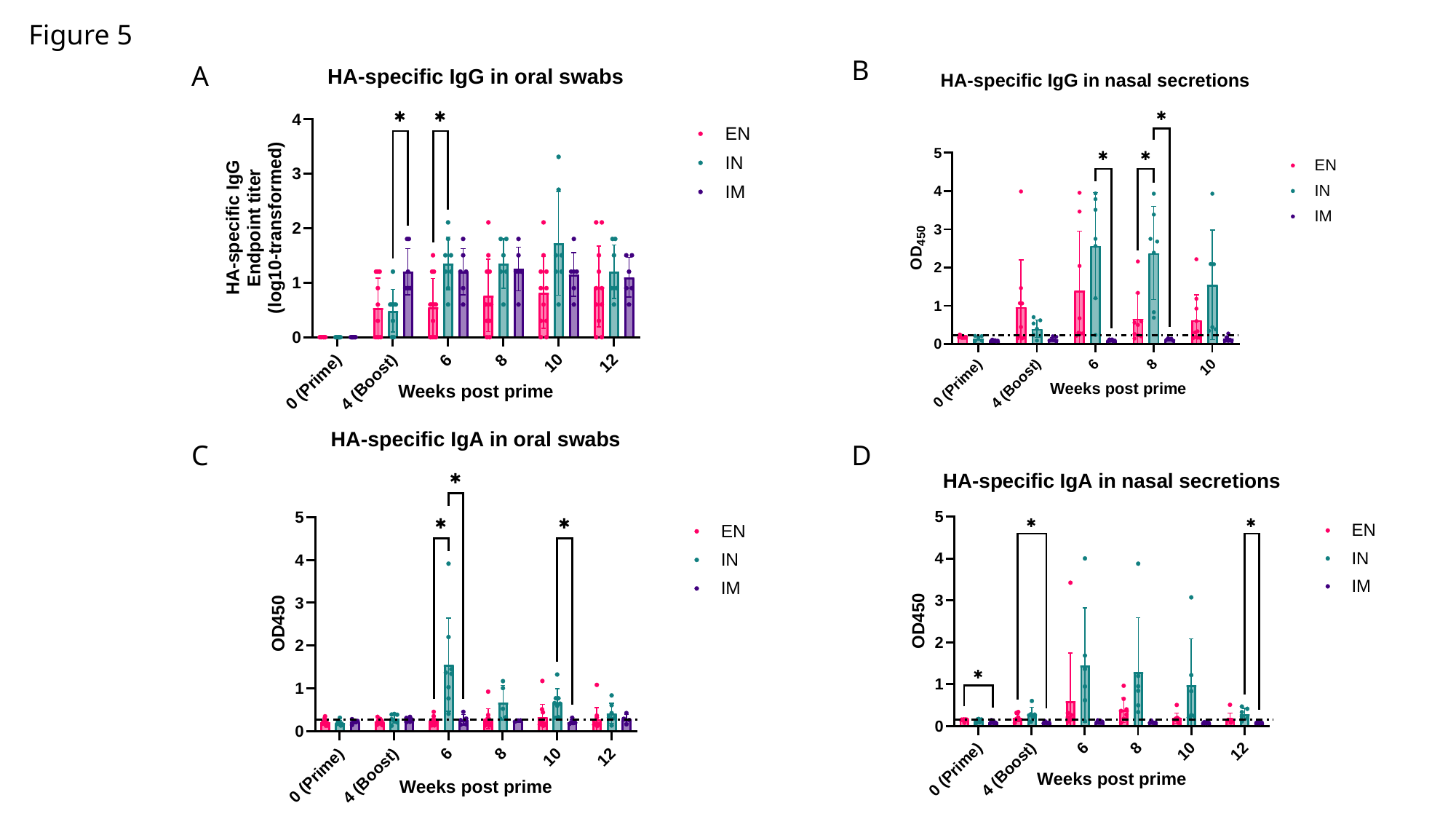

Figure 5
B
A
C
D

### Slide 8
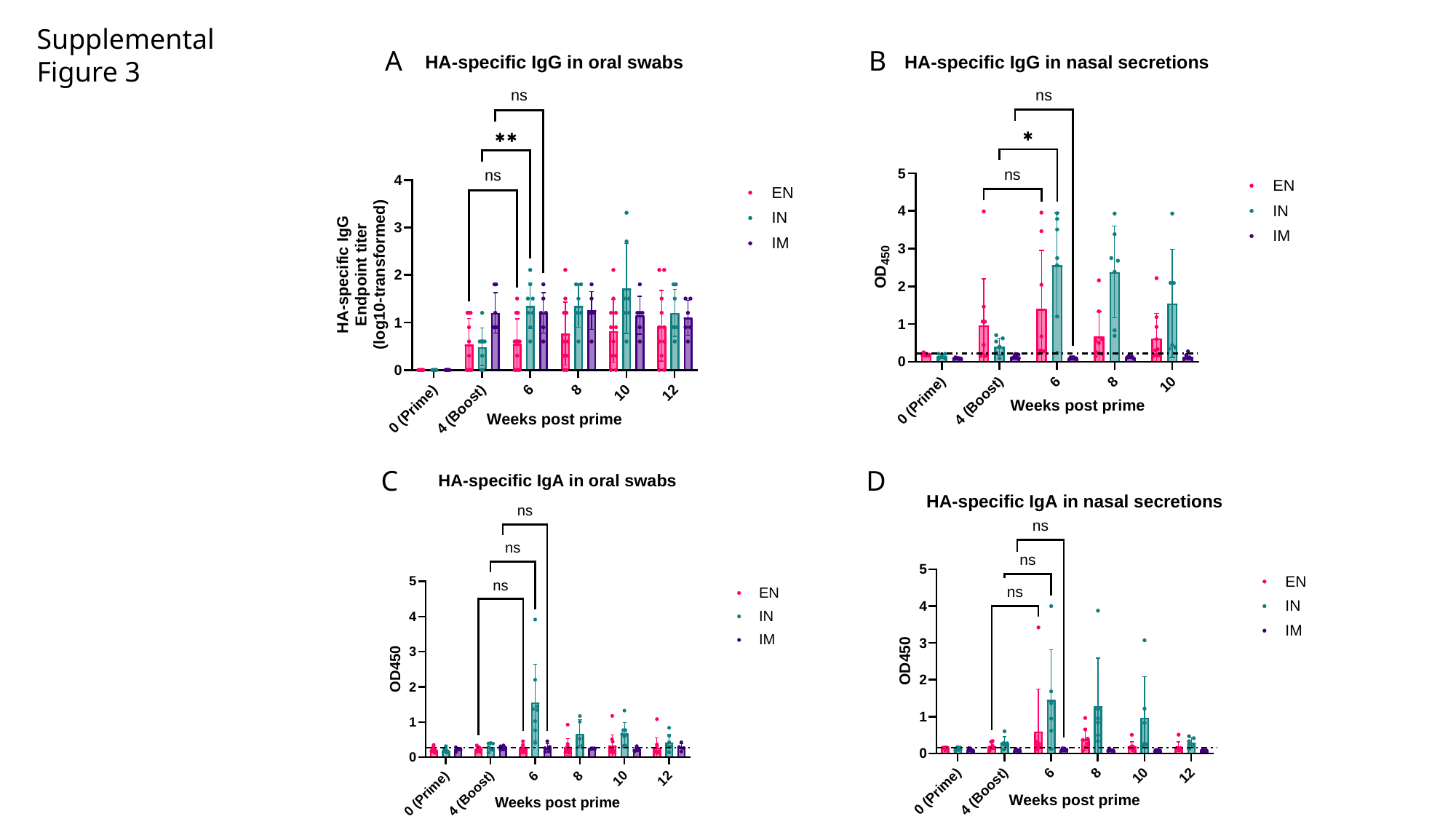

Supplemental Figure 3
A
B
C
D

### Slide 9
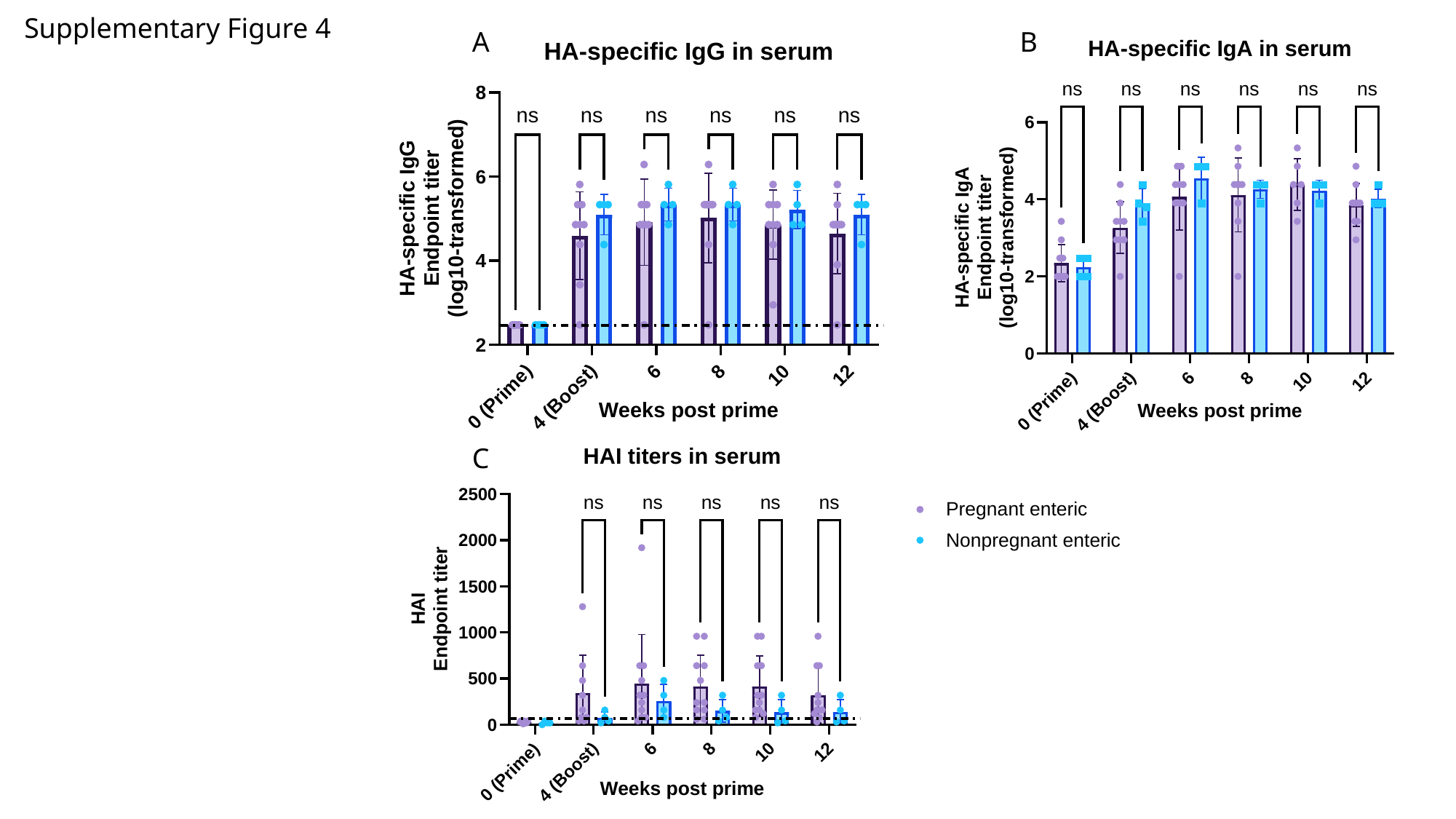

Supplementary Figure 4
A
B
C

### Slide 10
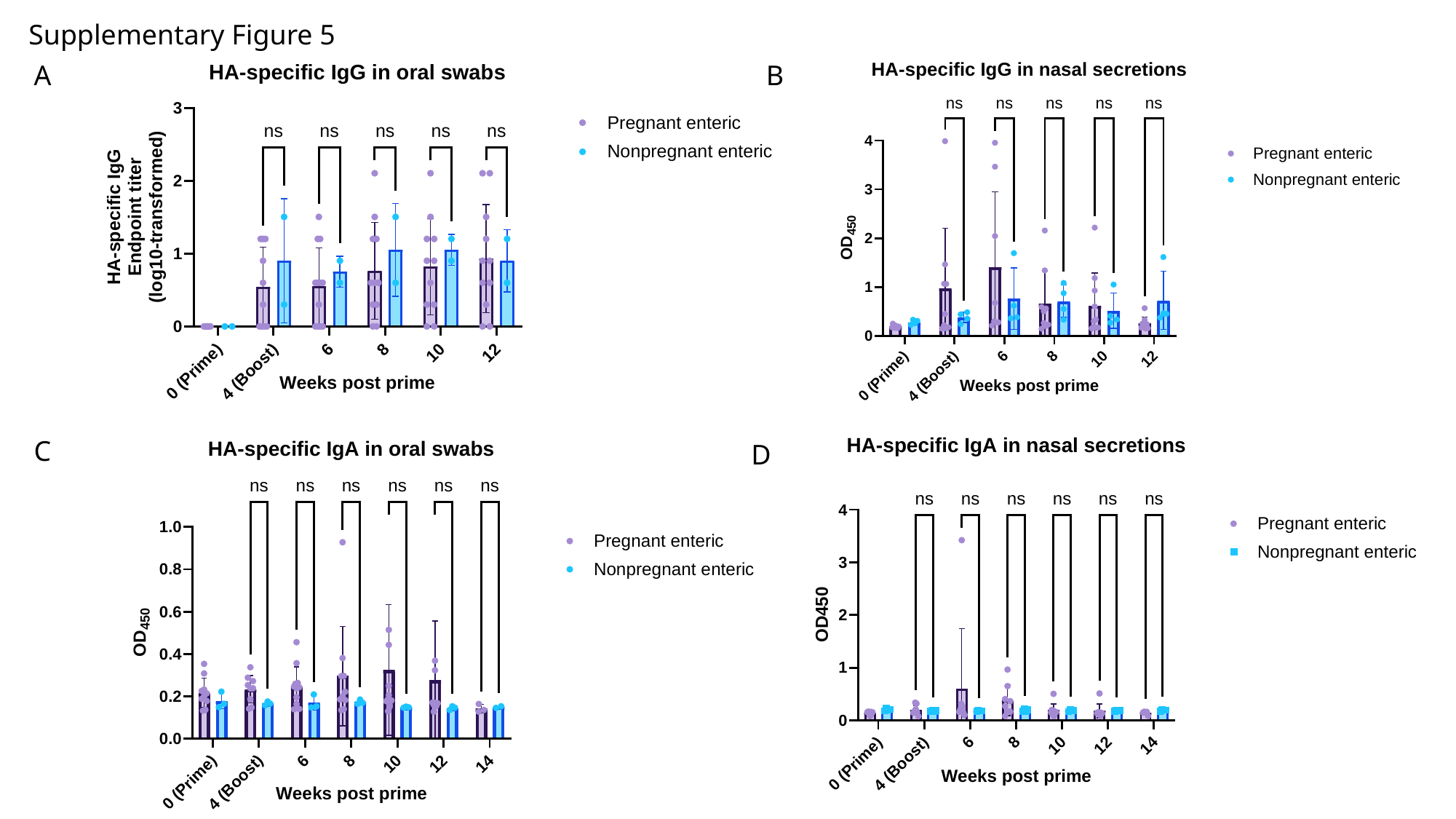

Supplementary Figure 5
A
B
C
D

### Slide 11
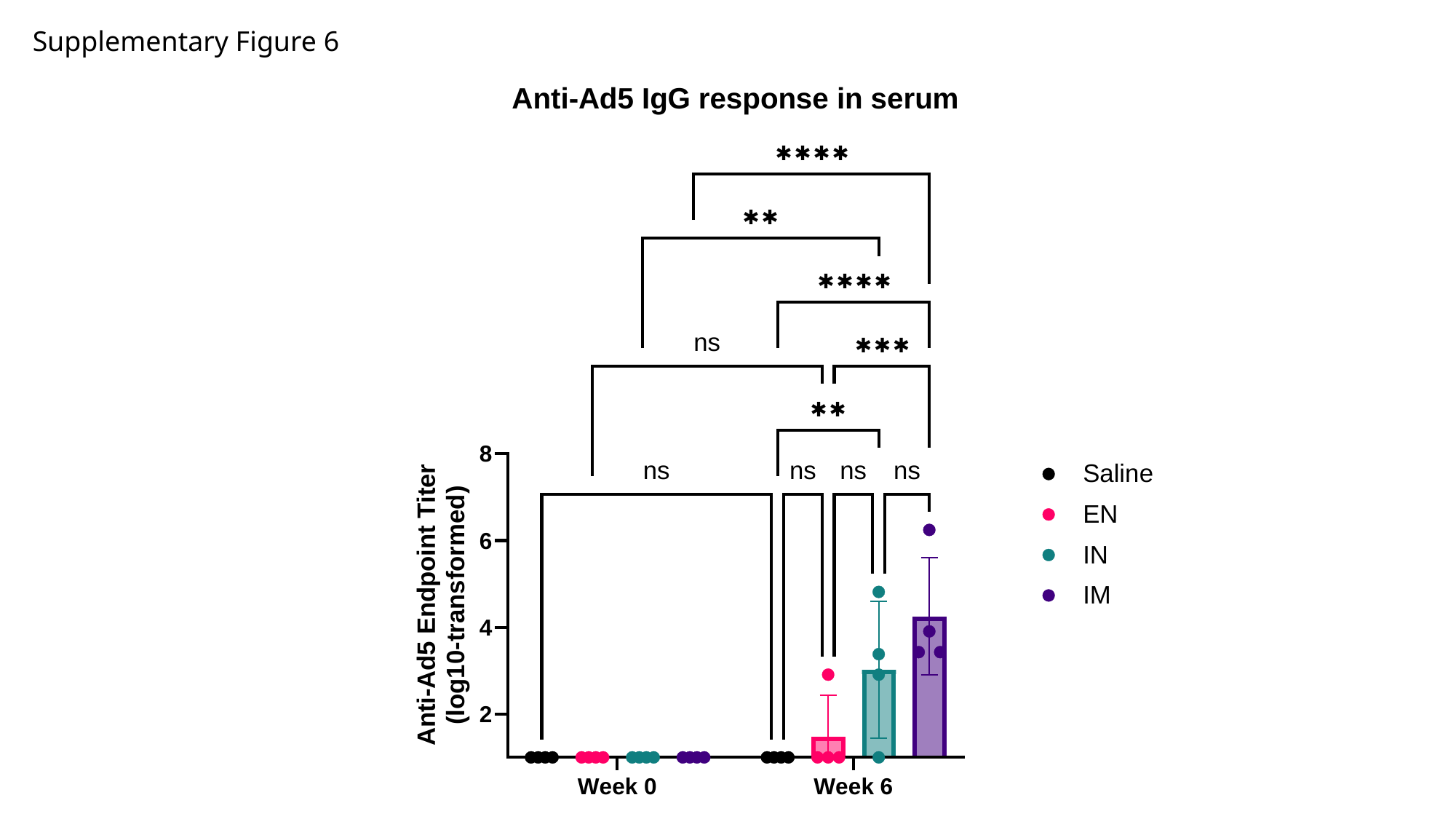

Supplementary Figure 6
